# q2-sample-classifier: machine-learning tools for microbiome classification and regression

**DOI:** 10.1101/306167

**Authors:** Nicholas A. Bokulich, Matthew Dillon, Evan Bolyen, Benjamin D. Kaehler, Gavin A. Huttley, J. Gregory Caporaso

## Abstract

Microbiome studies often aim to predict outcomes or differentiate samples based on their microbial compositions, tasks that can be efficiently performed by supervised learning methods. Here we present a benchmark comparison of supervised learning classifiers and regressors implemented in scikit-learn, a Python-based machine-learning library. We additionally present q2-sample-classifier, a plugin for the QIIME 2 microbiome bioinformatics framework, that facilitates application of the scikit-learn classifiers to microbiome data. *Random forest, extra trees*, and *gradient boosting* models demonstrate the highest performance for both supervised classification and regression of microbiome data. Automated feature selection and hyperparameter tuning enhance performance of most methods but may not be necessary under all circumstances. The q2-sample-classifier plugin makes these methods more accessible and interpretable to a broad audience of microbiologists, clinicians, and others who wish to utilize supervised learning methods for predicting sample characteristics based on microbiome composition. The q2-sample-classifier source code is available at https://github.com/qiime2/q2-sample-classifier. It is released under a BSD-3-Clause license, and is freely available including for commercial use.

## Introduction

A key goal of most microbiome studies is to discover patterns in microbial communities that relate to different groups of samples. For example, studies may test whether the microbiota differentiate patients receiving different treatments, or correlate with an environmental gradient such as pH or temperature. Microbiome sequencing data sets are often high-dimensional, and many experimental problems may benefit from the employment of machine learning methods for feature selection, pattern recognition, and prediction.

Supervised learning (SL) methods offer a powerful suite of tools for characterizing and differentiating microbial communities, with several promising applications for microbiome analysis (Knights, Kuczynski, *et al.*, 2011). The goal of SL is to train a machine learning model on a set of samples with known class labels, and then use that model to predict the class membership of additional, unlabeled samples. In the process, many learning models rank the importance of each feature to identify those that are most predictive of class membership. These class labels can either be categorical, e.g., a patient’s disease state or body site (a classification problem), or continuous numerical data, e.g., the age of a patient or concentration of a plasma metabolite (a regression problem). Sample classes may then be predicted for new, unknown samples, e.g., for the prediction of pathogen colonization outcomes (Schubert *et al.*, 2015) or wine metabolite abundance (Bokulich, Collins, *et al.*, 2016) based on baseline microbiome composition. The ability to categorize new samples, as opposed to describing the structure of existing data, is a major advantage of SL over conventional methods for microbiome analysis (Knights, Kuczynski, *et al.*, 2011). This characteristic extends itself to many useful applications, e.g., the prediction of disease/susceptibility (Yazdani *et al.*, 2016; Schubert *et al.*, 2015; Pasolli *et al.*, 2016), crop productivity (Chang *et al.*, 2017), sample collection site (Bokulich *et al.*, 2013), the identification of mislabeled samples in microbiome data sets (Knights, Kuczynski, *et al.*, 2011), or the detection of abnormal microbiota-for-age development in malnourished children (Subramanian *et al.*, 2014).

We describe q2-sample-classifier (https://github.com/qiime2/q2-sample-classifier), a QIIME 2 plugin (https://qiime2.org/) to support supervised learning tools for pattern recognition in microbiome data. This plugin is free and open-source software (BSD-3-Clause license), and currently implements several supervised learning methods, including regression and classification models that are benchmarked in the current study. This plugin accepts a feature table (sample X feature observations), consisting of frequency values (e.g., counts) for each feature (e.g., amplicon sequence variants or operational taxonomic units) observed in each sample. For many investigators using QIIME 2 and q2-sample-classifier, feature observations will most commonly consist of counts of amplicon sequence variants, operational taxonomic units, or taxa detected by marker-gene or shotgun metagenome sequencing methods, but other data, such as gene, transcript, protein, or metabolite abundance could be easily used as inputs.

## Results

We tested the performance of q2-sample-classifier using the following benchmark evaluations. First, we compared the relative performance of each estimator algorithm (described in detail in materials and methods) for sample prediction, using several well-characterized test data sets (Table 1). These are data from previous studies that exhibit distinct characteristics and a range of challenges to evaluate algorithm performance under different conditions. Second, we evaluated the influence of feature selection (FS) and hyperparameter tuning (HT) (see materials and methods) on method performance; each benchmark test was performed with FS and HT, with only FS or HT, or without any optimization.

**Table 1.**
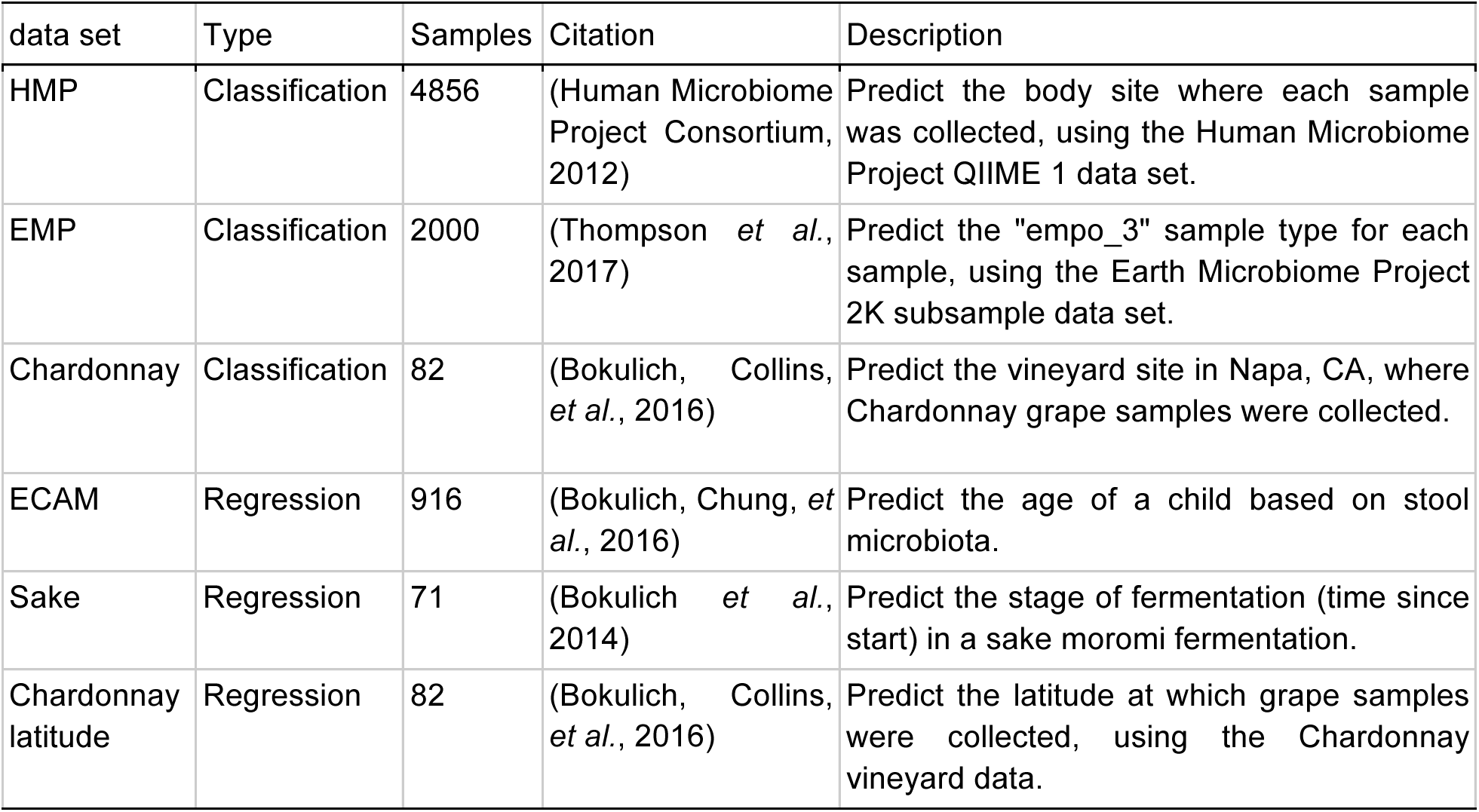
data sets employed for supervised learning benchmark tests.

### Sample classification

Classification tasks involve the prediction of categorical sample labels as a function of feature data. For example, a classifier may predict whether a sample is of a particular type (e.g., stool, soil), collected from a particular location (e.g., body sites, country of origin), or belongs to a healthy or diseased patient, based on that sample’s microbiota composition, metabolome, metagenome, genetic profile, or other feature data. q2-sample-classifier integrates several scikit-learn classification methods, enabling investigators to explore and answer these types of classification problems with their own data sets. We benchmarked the relative performance of classifiers on several different classification problems. To benchmark a range of classification problems, we predict the “empo_3” sample type of environmental samples in the Earth Microbiome Project (EMP) global survey (Thompson *et al.*, 2017); the vineyard of origin for Chardonnay grape samples collected from different vineyards in Napa County, California (Bokulich, Collins, *et al.*, 2016); and body sites in the Human Microbiome Project (HMP) 16S rRNA data (Human Microbiome Project Consortium, 2012). We note that these tests may not necessarily generalize to other classification tasks, and some aspects of classifier performance may depend on the nature of a given experiment, but these test data sets offer a useful range of different data characteristics for the purposes of our benchmark test. To illustrate the characteristics of these test data sets, we use principal coordinate analysis (PCoA) of UniFrac distance (Lozupone and Knight, 2005) to visualize the phylogenetic similarity among sample categories in these data sets (Fig. 1). Unweighted UniFrac distances relate to differences in sample composition, wherein differences indicate the presence of unique phylotypes; abundance-weighted UniFrac distances relate to differences in sample structure, or the relative abundance of each phylotype. The chardonnay vineyard data set is the smallest and simplest; some vineyards are easily distinguished visually on PCoA plots and show high intra-vineyard similarity while others exhibit higher spread and are more difficult to distinguish from each other (Fig 1A, B). The EMP data set contains a mixture of highly unique sample types (e.g., Non-saline sediment and plant rhizosphere), rough divisions of interrelated sample types (e.g., host-associated and non-host-associated sample types; and non-saline versus saline samples), and other sample types that exhibit a high degree of spread and are difficult to distinguish from other sample types (e.g., plant and animal corpus samples) (Fig 1C, D). The HMP data set roughly segregates into subclusters containing oral, skin, vaginal, and intestinal (stool) body sites, but within these subclusters samples are much more difficult to distinguish (Fig 1E, F).

**Figure 1.**
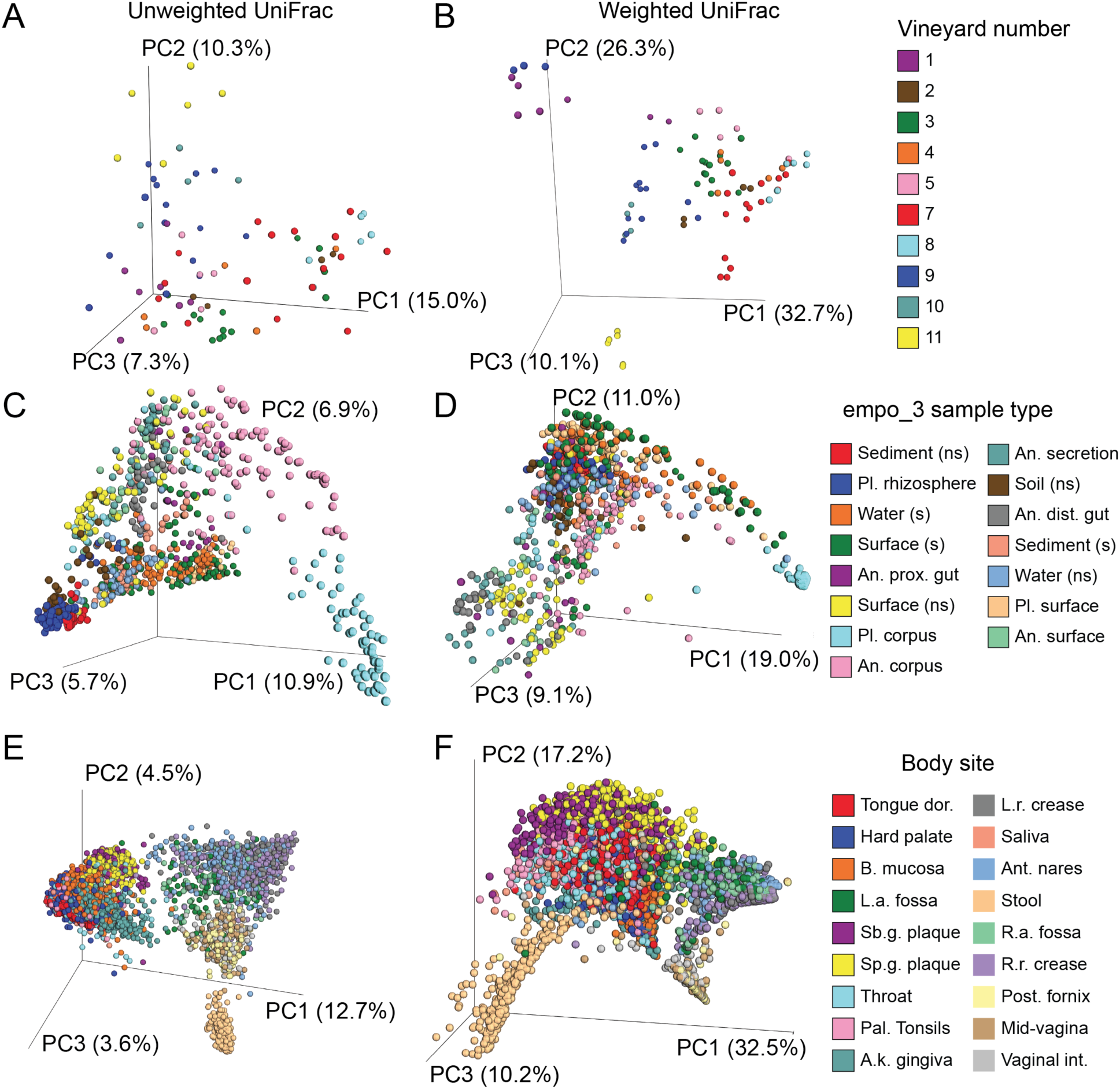
Principal coordinate analysis of chardonnay vineyard (A, B), EMP (C, D), and HMP (E, F) classification test data sets. Samples are colored by the categories used for classification. Ordinations are shown for unweighted (left: A, C, E) and weighted UniFrac (right: B, D, F). Abbreviations: Pl. = plant; s = saline; ns = non-saline; An. = animal; prox. = proximinal; dist. = distal; Tongue dor. = dorsum; B. mucosa = buccal; L.a./R.a. fossa = left/right anticubital; Sb.g./Sp.g. plaque = sub/supragingival; Pal. tonsils = palatine; A.k. gingiva = attached keratinized; L.r./R.r. crease = left/right retroauricular crease; Post. fornix = posterior; Vaginal int. = introitus.

Benchmarking results reveal common trends among classifier performance (Fig. 2). The best overall accuracy (percentage of test samples that were accurately classified) was achieved for the chardonnay vineyard data set (0.71–1.0), followed by EMP (0.65–0.86) and HMP (0.42– 0.65). However, this belies the strong performance of some classifiers for classifying EMP and HMP samples, which contained more classes and hence a lower baseline accuracy (i.e., classification accuracy if all samples were classified to the most abundant class). The accuracy ratio (overall / baseline) was highest for HMP classifiers (median 8.7; range 6.2–9.9), followed by EMP (6.2; 5.1–6.7) and chardonnay (5.0; 4.0–5.7). Inspection of confusion matrix heatmaps (Fig. 2) indicates that performance is dampened in the HMP data set by classification among closely related body sites: oral sites (throat, tongue, palatine tonsils, attached keratinized gingiva, buccal mucosa, hard palate, and to a lesser degree saliva) frequently cross-classify; antecubital fossae and retroauricular creases cannot be distinguished between left/right and frequently cross-classify as microbiologically similar skin sites; and vaginal sites (mid-vagina, vaginal introitus, and posterior fornix) commonly cross-classify. The EMP data set exhibits similar cross-classification issues among closely related empo_3 types, e.g., animal surface, secretion, and non-saline surfaces (Fig. 2).

**Figure 2.**
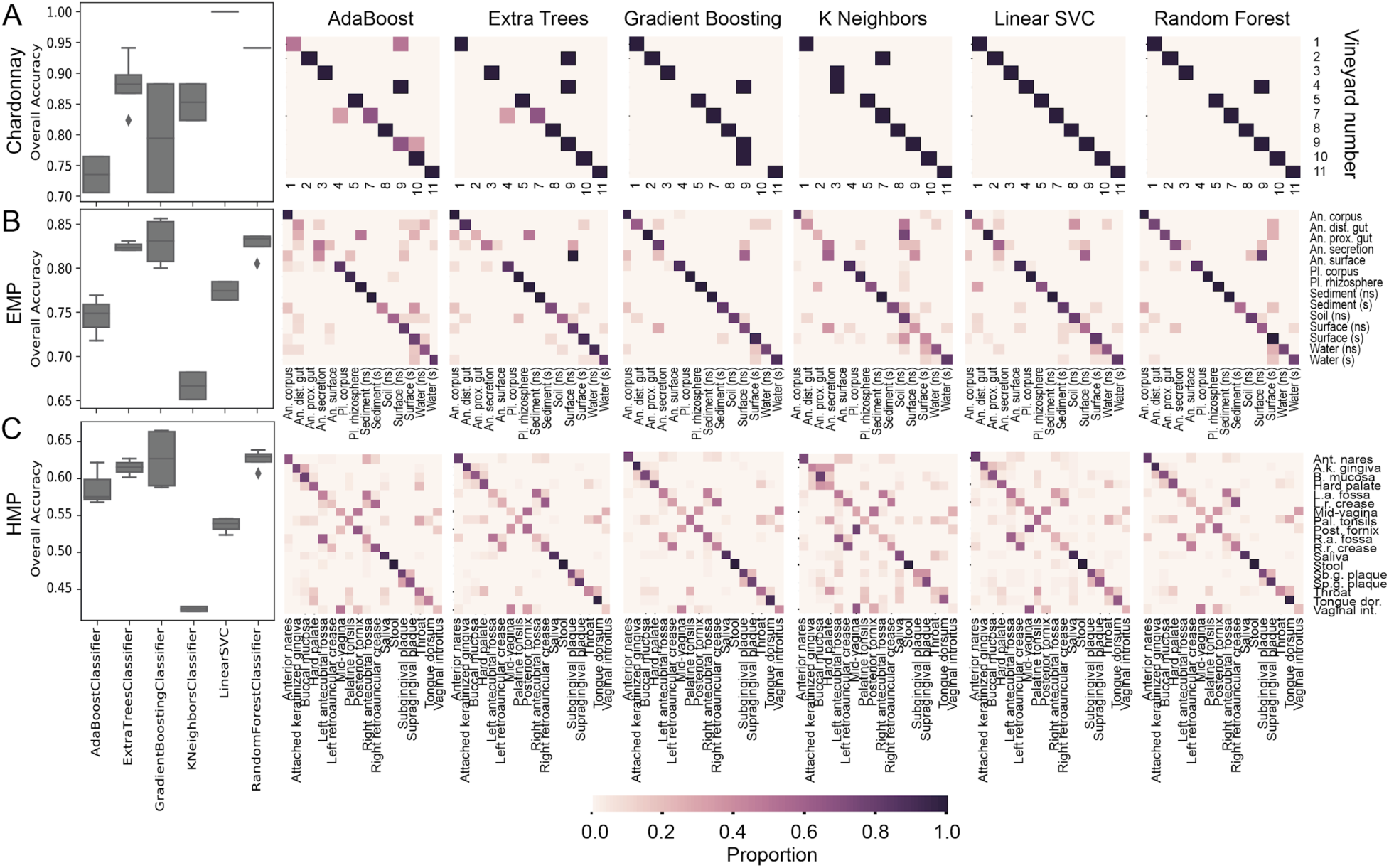
Classifier overall accuracy boxplots and confusion matrices for the chardonnay vineyard (A), EMP (B), and HMP (C) data sets. Boxplots show median and quartile accuracy for each classifier, averages across all optimization types (none, FS, HT, or both). Confusion matrices show the proportion of times each sample type was classified to each class by each classifier. Row labels indicate the true class and column labels indicate the predicted class and proportion (corresponding to the color key); correct classifications appear along a diagonal from upper-left to lower-right corners. Abbreviations: Pl. = plant; s = saline; ns = non-saline; An. = animal; prox. = proximinal; dist. = distal; Tongue dor. = dorsum; B. mucosa = buccal; L.a./R.a. fossa = left/right anticubital; Sb.g./Sp.g. plaque = sub/supragingival; Pal. tonsils = palatine; A.k. gingiva = attached keratinized; L.r./R.r. crease = left/right retroauricular crease; Post. fornix = posterior; Vaginal int. = introitus.

The ensemble methods *random forest* (0.83 median overall accuracy), *extra trees* (0.82), and *gradient boosting* (0.75) classifiers performed best across all three data sets (Table 2). For the chardonnay vineyard data set, *linear SVC* achieved the best overall accuracy (100% of test samples were accurately classified), but this method achieved mediocre performance with the other two data sets (Fig. 2A). This likely relates to the unique characteristics of the chardonnay data set, particularly the relative simplicity of this classification problem (e.g., compared to differentiating left/right antecubital fossae in the HMP data set); *K neighbors* classifiers also performed relatively well on this data set though this method achieved the worst performance on the other two data sets (0.67 median overall accuracy).

**Table 2.** Median classifier accuracy, accuracy ratio, and improvements in overall accuracy from hyperparameter tuning (HT) and feature selection (HS), averaged across test data sets.

| Estimator | Overall Accuracy | Accuracy Ratio | HT median change | FS median change | HT+FS median change |
| --- | --- | --- | --- | --- | --- |
| **AdaBoost** | 0.71 | 5.72 | -0.05 | 0.06 | -0.02 |
| **ExtraTrees** | 0.82 | 6.42 | -0.01 | 0.00 | 0.01 |
| **GradientBoosting** | 0.75 | 6.48 | 0.07 | 0.00 | 0.08 |
| **KNeighbors** | 0.67 | 5.20 | 0.03 | 0.00 | 0.03 |
| **LinearSVC** | 0.77 | 6.04 | 0.00 | 0.01 | 0.02 |
| **RandomForest** | 0.83 | 6.50 | 0.00 | 0.00 | 0.00 |

FS and HT can be important aspects of supervised learning model optimization. We simultaneously benchmarked the effects of automatic FS and HT steps implemented in q2-sample-classifier on classifier performance (Table 2, Figure 2-3). The sensitivity of each classification method to parameter optimization can be observed in the boxplot quartile measurements shown in Figure 2; The magnitude of accuracy improvement can be observed in heatmaps of model performance (Figure 3) and in Table 3. *Random forest* classifier accuracy was mostly unaffected by HT and FS steps (0.0 median accuracy change), whereas *gradient boosting* (0.07) and *k-neighbors* classifiers (0.03) were highly affected by HT. Only *AdaBoost* (0.06) saw a dramatic change due to FS, suggesting that FS may be unnecessary for many models that are not re-used with other data sets. FS involves repeatedly training and testing the model, increasing computational runtime; however, a sparser model that uses fewer features will require less runtime to classify new samples, and hence models that are re-used repeatedly will still benefit from FS even if accuracy is not markedly improved. FS can also be an end in itself: identifying a minimal set of features that predicts a class/value with maximal accuracy can be valuable for selecting targets for downstream study and establishment of biomarkers. Feature importance is reported by the classification and regression methods implemented in q2-sample-classifier, facilitating the use of this information for downstream analyses.

**Figure 3.**
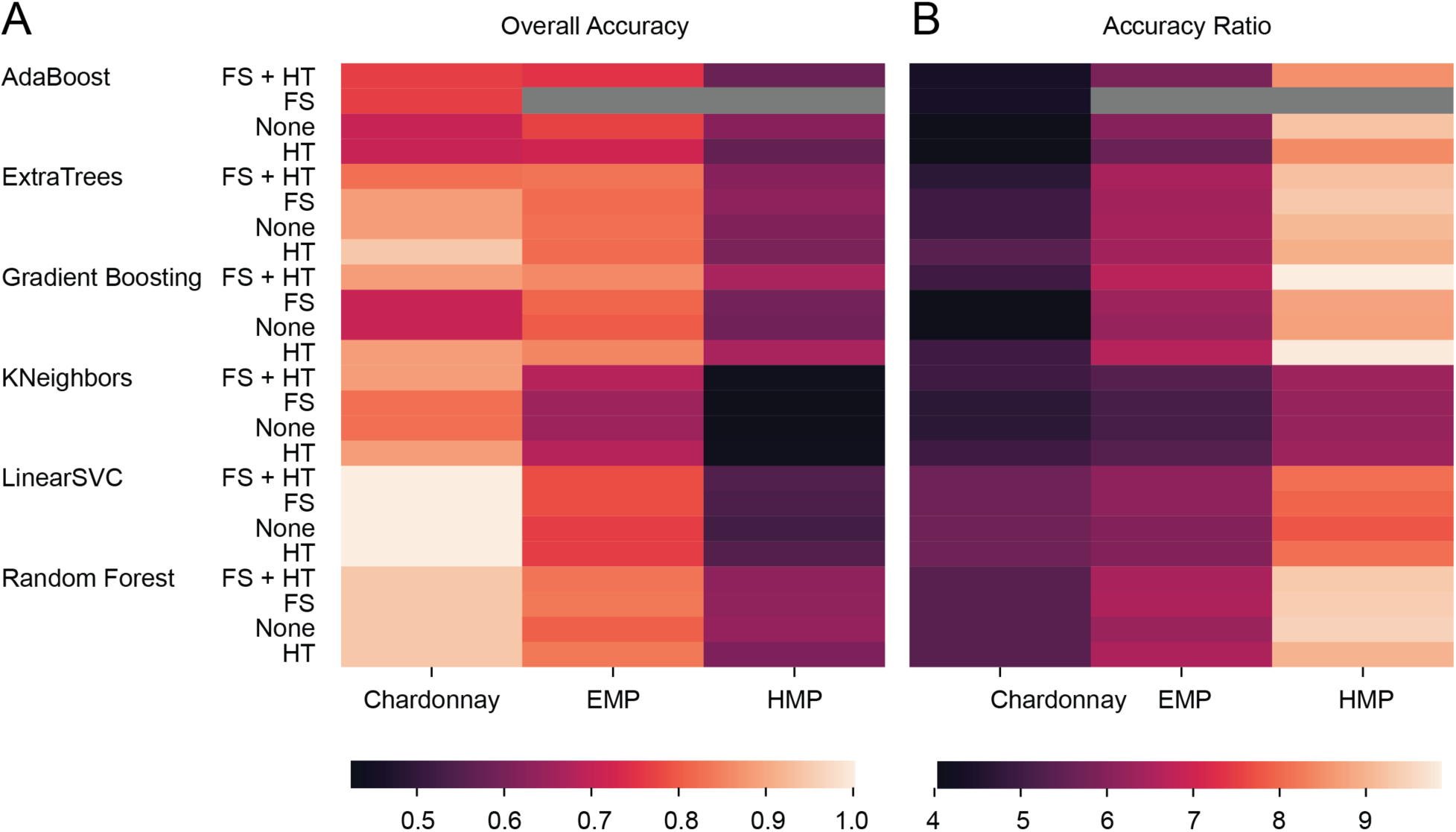
Heatmap of overall classification accuracy (A) and overall / baseline accuracy ratio (B) for each estimator, optimization method (none, FS, HT, or both), and test data set. Grey boxes were not computed.

### Sample regression

Regressors predict a numerical target value as a function of feature data. For example, a regressor may predict the abundance of a metabolite, sample pH, or the geospatial coordinates where a sample was collected. q2-sample-classifier supports several scikit-learn regression methods. We benchmarked the relative performance of regressors for predicting the age of children as a function of their stool microbiota in the early childhood and the microbiome (ECAM) study (Bokulich, Chung, *et al.*, 2016), the age of sake fermentations as a function of their bacterial composition (Sake study; (Bokulich *et al.*, 2014)), and the geocoordinates of chardonnay grape samples as a function of their bacterial compositions (chardonnay latitude (Bokulich, Collins, *et al.*, 2016)) (Table 1). These data sets exhibit different sample ordination characteristics (Fig. 4). The ECAM data set is characterized by a gradual transition from birth to older childhood by both weighted and unweighted UniFrac PCoA (Fig. 4A, B). Sake samples cannot be distinguished by unweighted UniFrac (Fig. 4C), but weighted UniFrac PCoA shows a shift from highly disparate microbial structure at the start of fermentation to tighter similarity among samples as fermentation progresses (Fig. 4D). The latitude of chardonnay samples show some clustering patterns by both UniFrac metrics, but a relationship between ordination and latitude cannot be discerned from PCoA (Fig. 4E, F).

**Figure 4.**
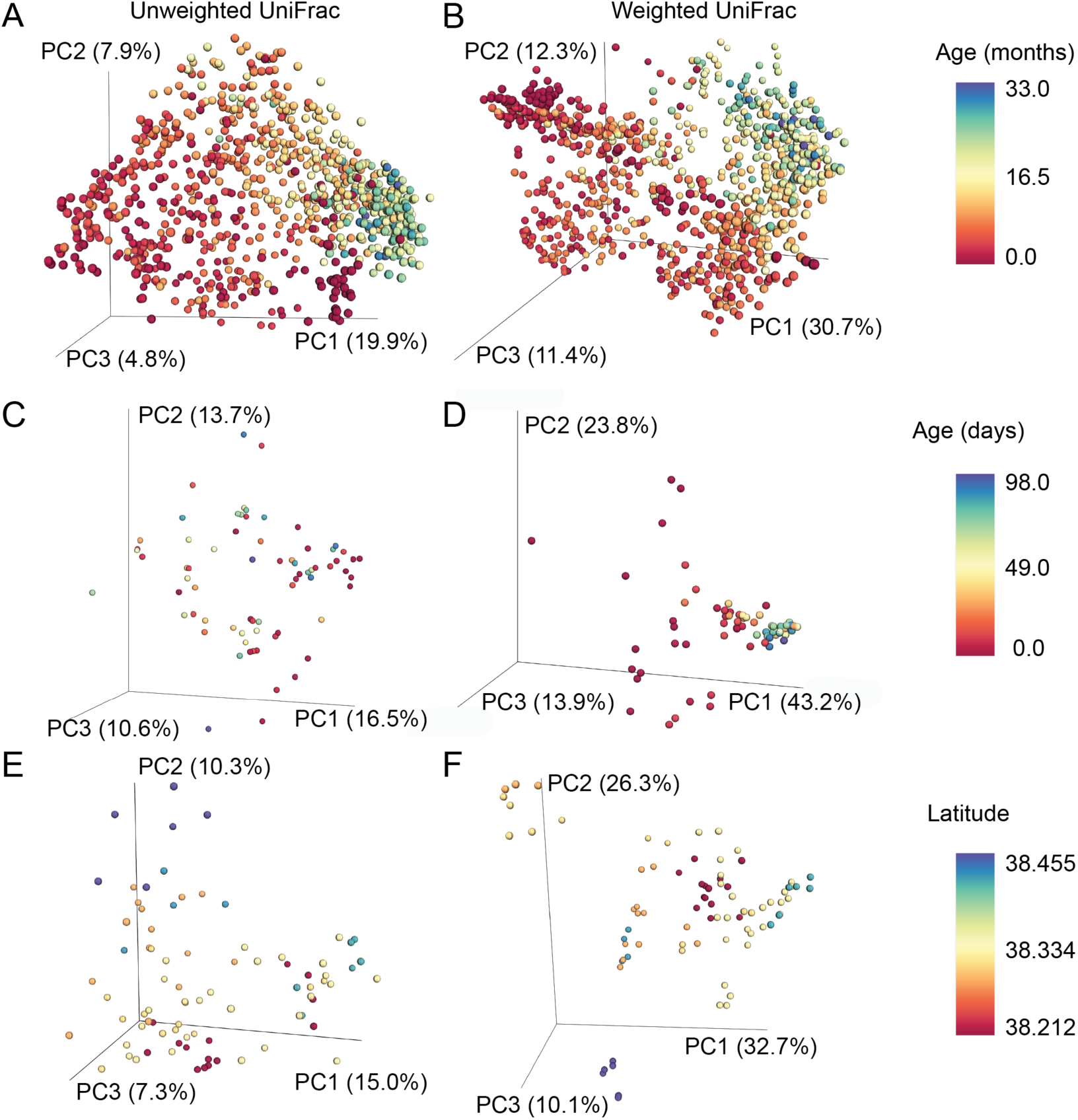
Principal coordinate analysis of ECAM (A, B), sake (C, D), and chardonnay (E, F) supervised regression test data sets. Samples are colored by the continuous sample data used for classification. Ordinations are shown for unweighted (left: A, C, E) and weighted UniFrac (right: B, D, F).

Regression accuracy is assessed based on the correlation between the true and predicted values for each sample; the mean square error (MSE) or linear least-squares regression coefficient (R) indicates the relative degree of accuracy (lower MSE and higher R values indicate better predictive performance). As we see a wide variance in MSE values both between and within data sets, we take the log MSE normalized within each data set *(log MSE – min max)* (nlMSE) for visualization and comparison purposes here.

Across all data sets, the best regression predictive accuracies were achieved with *gradient boosting* (0.16 median nlMSE), *extra trees* (0.17), and *random forest* (0.18) ensemble regressors (Table 3, Fig. 5). *Lasso* (0.68), *ElasticNet* (0.74), *Ridge* (0.78), and *linear SVR* regressors (0.89) demonstrate universally poor performance. The performance of *k-neighbors* regressors (0.31) appears to vary widely by data set and optimization; FS+HT optimized *k-neighbors* regressors achieve the lowest nlMSE score among all regressors for the chardonnay latitude data set (Fig. 5A).

**Figure 5.**
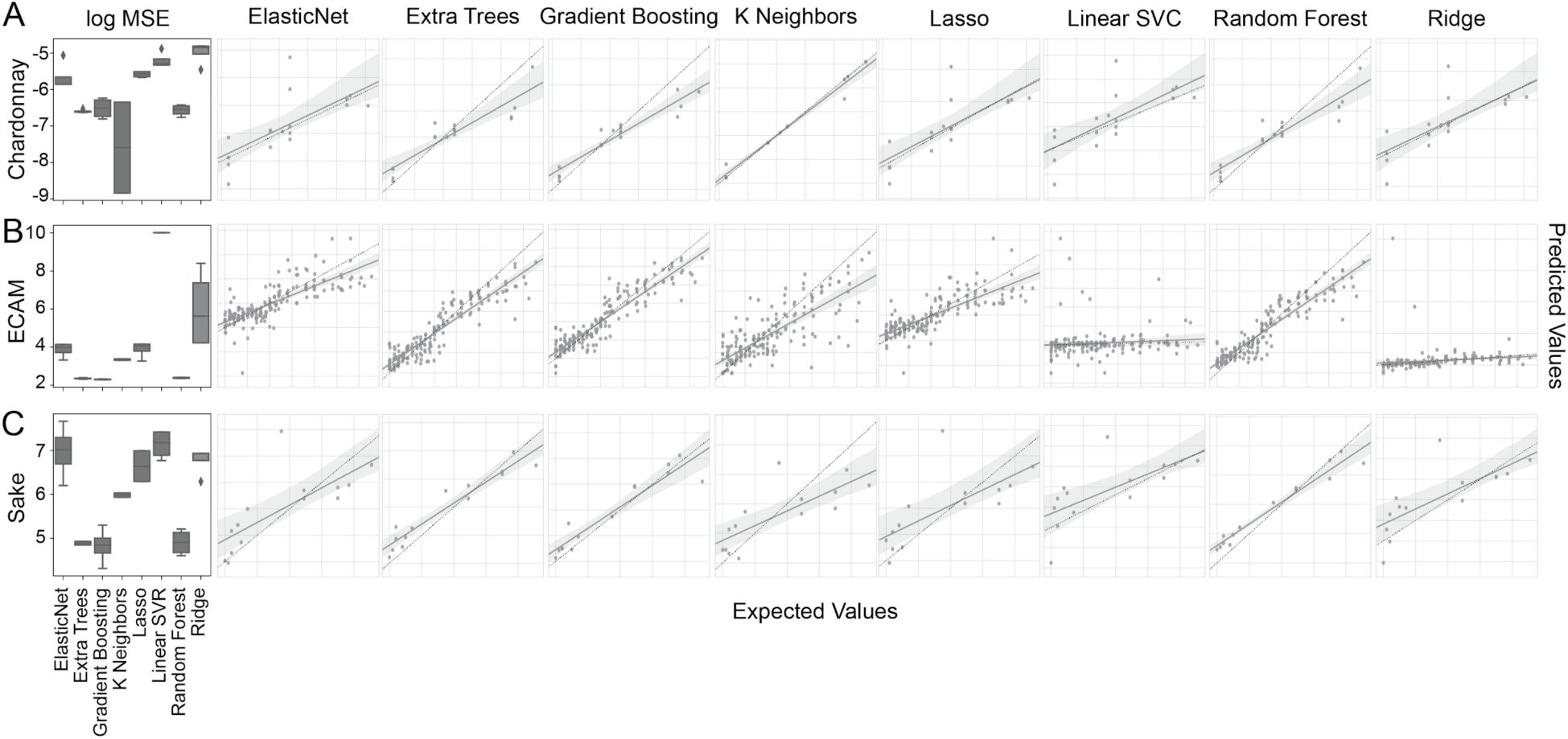
Supervised regression model normalized log mean square error (nlMSE) boxplots and regression plots for prediction of chardonnay vineyard latitudes (A), children’s ages in the ECAM study (B), and the age (days) of sake fermentations (C). Boxplots show median and quartile nlMSE for each regressor, averages across all optimization types (none, FS, HT, or both). Regression plots show the correlation between true values (x-axes) and predicted values (y-axes) for each sample. A linear regression fit and confidence intervals are shown as a dark grey line and light grey shading. A dashed line indicates the true 1:1 ratio between axes.

**Table 3.** Median regressor mean square error (MSE), linear least-squares regression coefficient (R), normalized log MSE (nlMSE), and nlMSE improvement from hyperparameter tuning (HT) and feature selection (HS), averaged across test data sets.

| Estimator | MSE | R | normalized MSE | HT median change | FS median change | HT+FS median change |
| --- | --- | --- | --- | --- | --- | --- |
| **ElasticNet** | 54.89 | 0.71 | 0.74 | -0.07 | 0.04 | 0.04 |
| **ExtraTrees** | 10.54 | 0.93 | 0.17 | 0.00 | 0.01 | 0.00 |
| **GradientBoosting** | 10.00 | 0.93 | 0.16 | -0.12 | -0.02 | -0.14 |
| **KNeighbors** | 28.39 | 0.79 | 0.31 | -0.01 | 0.00 | -0.01 |
| **Lasso** | 58.37 | 0.65 | 0.69 | -0.09 | 0.00 | -0.03 |
| **LinearSVR** | 948.13 | 0.49 | 0.89 | -0.05 | 0.07 | 0.08 |
| **RandomForest** | 10.93 | 0.93 | 0.18 | -0.05 | -0.01 | -0.08 |
| **Ridge** | 304.24 | 0.42 | 0.78 | -0.18 | 0.00 | 0.00 |

Model optimization evidently has a more substantial impact on supervised learning regressors compared to classifiers (Table 3, Fig. 5-6). Automatic HT appears to have a beneficial impact on all regressors (median change in nlMSE range -0.18 to -0.01) except for *extra trees* (0.0). Automatic FS has a negligible effect on most estimators (but may still be beneficial, as discussed above) and negatively impacts *ElasticNet* (0.04) and *linear SVR* regressors (0.07). This negative impact occurs because FS can conflict with the L1 and/or L2 normalization that these models already employ for sparse feature regression, resulting in excessive feature elimination.

**Figure 6.**
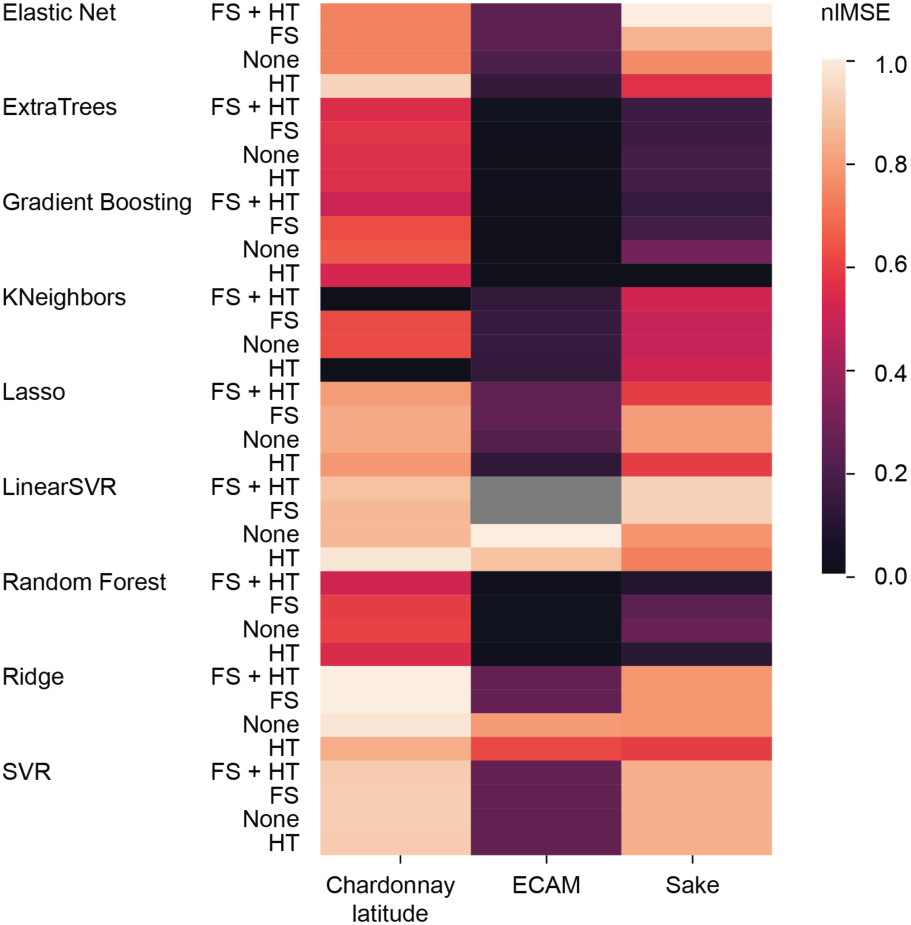
Heatmap of regression accuracy normalized log mean square error (nlMSE) for each estimator, optimization method (none, FS, HT, or both), and test data set. Lower nlMSE is better, and nlMSE is normalized across columns (data sets) such that 0.0 = the best-performing method and 1.0 = the worst-performing method for that data set. Grey boxes were not computed.

## Conclusions

Here we present a microbiome-focused benchmark of machine learning classifiers and regressors implemented in scikit-learn. *Random forest, extra trees*, and *gradient boosting* models appear to perform best for both classification and regression of microbiome data. *K-neighbors* and *linear support vector* machines methods may be faster, accurate substitutes for easily classifiable data sets (e.g., the chardonnay vineyard test data employed here), but perform poorly on more complex microbial data sets. Automated method optimization, as implemented in q2-sample-classifier, boosts performance of most methods but may not be necessary under all circumstances. Implementation in q2-sample-classifier will make these methods more accessible and interpretable to a broad audience of microbiologists, clinicians, and others who wish to utilize supervised learning methods for predicting sample characteristics based on microbiome composition.

## Materials and methods

### Supervised learning models and plugin design

The q2-sample-classifier plugin is accessible by multiple user interfaces supported in QIIME 2 (Caporaso *et al.*, 2010); a Python 3 application programming interface (API) is accessible for advanced users, and non-programmers may access this plugin via QIIME 2’s command-line interface (CLI) or q2studio’s graphical user interface (GUI). Here we describe and benchmark the classify-samples and regress-samples actions that are exposed in the CLI/GUI (API users have access to the same functionality). These functions perform identical steps but access different estimators (classification and regression models, respectively) and produce different visualizations relevant to these tasks. The q2-sample-classifier plugin is written in Python 3.5 and employs pandas (McKinney, 2010) and numpy (van der Walt *et al.*, 2011) for data manipulation, scikit-learn (Pedregosa *et al.*, 2011), and scipy (https://www.scipy.org/) for statistical testing, and matplotlib (Hunter, 2007) and seaborn (https://zenodo.org/record/12710) for data visualization. Plugin method descriptions, API, and tutorials can be accessed at https://docs.qiime2.org/.

All q2-sample-classifier actions accept a feature table (e.g., of amplicon sequence variant, OTU, taxon, gene, transcript, or metabolite abundance data) and sample metadata file as input, and require a metadata column name to use as a prediction target. Input samples are shuffled and split into training and test sets at a user-defined ratio (default: 4:1) with or without stratification (stratified by default); test samples are left out of all model training steps and are only used for final model validation.

The plugin incorporates various methods available in scikit-learn (Pedregosa *et al.*, 2011) to support a range of common supervised learning algorithms. The following estimators are currently implemented in q2-sample-classifier and tested here: *AdaBoost* (Freund and Schapire, 1997), *Extra Trees* (Geurts *et al.*, 2006), *Gradient boosting* (Friedman, 2002), and *Random Forest* (Breiman, 2001) ensemble classifiers and regressors; *linear SVC, linear SVR*, and *non-linear SVR* support vector machine classifiers/regressors (Cortes and Vapnik, 1995); and *k-Neighbors* classifiers/regressors (Altman, 1992). In addition, the following estimators can be used for sample regression: *Elastic Net* (Zou and Hastie, 2005), *Ridge* (Hoerl and Kennard, 1970), and *Lasso* (Tibshirani, 1996) regression models.

Feature selection and hyperparameter tuning are performed automatically and optionally. The user can toggle these functions on/off, and can select the number of cross-validations to perform for each (default = 5). Feature selection is performed using cross-validated recursive feature elimination (http://scikit-learn.org/stable/modules/generated/sklearn.feature_selection.RFECV.html) to select the features that maximize predictive accuracy. Hyperparameter tuning is automatically performed using a cross-validated randomized parameter grid search (http://scikit-learn.org/stable/modules/generated/sklearn.model_selection.RandomizedSearchCV.html) to find hyperparameter permutations (within a sensible range) that maximize accuracy.

Training data sets are then used to fit prediction models with optimized hyperparameters and features. Fitted models then predict the classes/values of the test data to evaluate model accuracy. The classify-samples and regress-samples actions that are exposed in the CLI/GUI create visualization files that include summaries of model accuracy and hyperparameters; confusion matrices (for classification) or linear regression scatter plots (for regression) of predicted and expected classes/values for each test sample; recursive feature selection; and feature importances.

### Test data sets

All test data sets are derived from previous studies, as described in Table 1. Data accessibility is described in the original studies. Feature tables from these studies all consisted of 16S rRNA gene sequencing operational taxonomic unit data. Microbial feature tables containing operational taxonomic unit or amplicon sequence variant observations (as defined by the original studies) were imported into QIIME 2, which was used to compute UniFrac (Lozupone and Knight, 2005) distance matrices and principal coordinates via the q2-diversity plugin (https://github.com/qiime2/q2-diversity), and plot PCoA plots via the q2-emperor plugin (Vázquez-Baeza *et al.*, 2013).

### Benchmark evaluation

Each estimator was used for classification/regression of the appropriate test data sets with full parameter optimization; with only hyperparameter tuning (HT) or feature selection (FS); or without any optimization. Each sample was subsampled so that 80% of samples were used for model training and 20% were held out as a test set for determining the predictive accuracy of each trained model. Whenever parameter optimization was enabled, HT and FS were optimized via 5-fold cross-validation of the training set. Model accuracy is reported as either the percentage of test samples that were predicted to belong to the correct class (for classifiers), or mean square error (for regressors).

## Acknowledgments

The authors thank Jai Ram Rideout for his input and assistance integrating q2-sample-classifier into QIIME 2. This project was funded in part by NSF Award 1565100 to JGC, and by the Partnership for Native American Cancer Prevention (NIH/NCI U54CA143924 and U54CA143925) to JGC.

